# Molecular Analyses Place the Genus *Keraunea* Outside Convolvulaceae

**DOI:** 10.1101/2022.11.14.516456

**Authors:** Pablo Muñoz-Rodríguez, John R.I. Wood, Lucía Villaescusa González, Charles C. Davis, Zoë A. Goodwin, Robert W. Scotland

## Abstract

The genus *Keraunea* was recently described in the Convolvulaceae Juss. family. Two species are currently recognised, both from Brazil. Molecular sequence data using three commonly applied DNA markers (*matK, rbcL* and the *nuclear ribosomal Internal Transcribed Spacer*) show that neither species is correctly placed in Convolvulaceae but indicates that the type, *K. brasiliensis*, should be placed in Malpighiaceae despite several morphological anomalies. The second species, *K. capixaba*, should be placed in Ehretiaceae. Given the surprising nature of these results, further studies are recommended before formal reclassification of these two taxa is made.

## TEXT

*Keraunea* Cheek & Sim.-Bianch. is a plant genus endemic to Brazil initially described in the family Convolvulaceae [1]. The genus includes two accepted species: *K. brasiliensis* Cheek & Sim.-Bianch., the type species described from the Brazilian state of Bahía with a paratype from Minas Gerais [1], and *K. capixaba* Lombardi described from Espirito Santo [2]. A few additional specimens have subsequently been ascribed to these two species, all of them from the same states where the species were first collected (see, e.g., records in GBIF: https://doi.org/10.15468/dl.e9wrys). In this note, we report the results of our study of images and herbarium specimens together with phylogenetic analyses of DNA sequence data sampled from type specimens of both species, which demonstrate the genus is misplaced in Convolvulaceae.

As part of our ongoing investigations of American Convolvulaceae [3–7], we came across high-resolution images of the type specimens of both *Keraunea brasiliensis* (K000979156) and *K. capixaba* (SP003725) available via JSTOR plants, as well as images of additional specimens available via the Reflora portal (http://reflora.jbrj.gov.br/reflora/herbarioVirtual). These specimens immediately attracted our attention because they did not resemble any American Convolvulaceae we have studied to date. The drawings and photographs in the original publications [1,2] are not an especially good fit with the morphology of Convolvulaceae, and there are some discrepancies between our observations of the types and the original descriptions and illustrations. Moreover, as indicated in the original publication [1], *Keraunea brasiliensis* resembles the Convolvulaceae genus *Neuropeltis* Wall., but *Neuropeltis* is restricted to the Palaeotropics, with a disjunct distribution in East Tropical Africa (c. 9 species) and South East Asia and India (4 species) [8,9].

To further explore this question, we studied all *Keraunea brasiliensis* collections cited in the original publication: *L. Passos 5263* (isotype K000979156), *J.A. Lombardi 1819* and *J.A. Lombardi 2107*. We studied these collections directly at the Kew herbarium or via high-resolution images in virtual herbaria. Similarly, we studied three *K. capixaba* collections listed in the original publication: *G.S. Siqueira 891* (isotype SP476897), *D.A. Folli 7117* and *G.S. Siqueira 893*. To the best of our knowledge, the *Keraunea* species had not yet been sequenced when we began our study, or the sequences had not been made available [cf. 10]. Thus, in addition to our morphological studies, we also sampled and sequenced three of these collections to incorporate them in the Convolvulaceae phylogenies generated as part of our ongoing systematic studies of the family [5].

We sequenced two *Keraunea brasiliensis* specimens (the isotype *Passos 5263* and the paratype *Lombardi 1819*) and one *K. capixaba* specimen (the paratype *G.S. Siqueira, 893*). We sequenced three DNA markers: the *nuclear ribosomal Internal Transcribed Spacer (nrITS) (Passos 5263* and *Lombardi 1819*), and the chloroplast *matK* (all specimens) and *rbcL* regions (*Passos 5263*). We extracted total genomic DNA using the Qiagen DNEasy extraction kit, and used primers AB101 and AB102 for *nrITS* amplification [11]; 413f-1 and 1227r-3 for *matK* amplification [12]; and rbcL-1F and rbcL-1460R for *rbcL* amplification [13,14]. In all cases, we used a reagent volume of 15 μl (7.3 μl H2O, 3 μl buffer, 0.7 μl MgCl2, 0.3 μl of each primer, 0.5 μl dNTPs, 1 μl BSA, 0.4 μl Taq polymerase, 1.5 μl sample DNA) for PCR amplification and standard PCR conditions (5’ at 80°C; 30 cycles of 1’ at 95°C, 1’ at 50°C, and 4’ at 65°C and a final stage of 4’ at 65°C). We cleaned the PCR reactions using the GeneJET PCR purification kit. We sequenced the samples using Sanger sequencing at Source Biosciences in Cambridge, United Kingdom, with the same primers used in the PCR. Furthermore, in light of our results (see below) we repeated both DNA extractions and amplifications in a different laboratory, with identical results.

To place the *Keraunea* samples in a phylogenetic context, we first queried all our sequences against the NCBI database using BLAST [15]. We also inferred a *matK* Angiosperm phylogeny using DNA sequence data previously published [16,17]. We aligned the sequences using MAFFT v.7.310 (Katoh & Standley 2013, 2016) and used Geneious v.9.1.8 to remove all columns in the alignments with 90% or more gaps. We then inferred a Maximum Likelihood phylogeny using IQ-Tree (Nguyen *et al*. 2015) with automatic model selection using ModelFinder (Kalyaanamoorthy *et al*. 2017) and 1,000 bootstrap replicates (-czb -bb 1000 -alrt 1000). The substitution model, TVM+F+I+G4, was selected based on the Bayesian Information Criterion. In the resulting phylogeny (Figure 1), we collapsed all nodes with less than 50% bootstrap support (-minsup 0.5). All subsequent phylogenies inferred in this study followed the same methodology, with substitution models indicated in each case.

**Figure 1.**
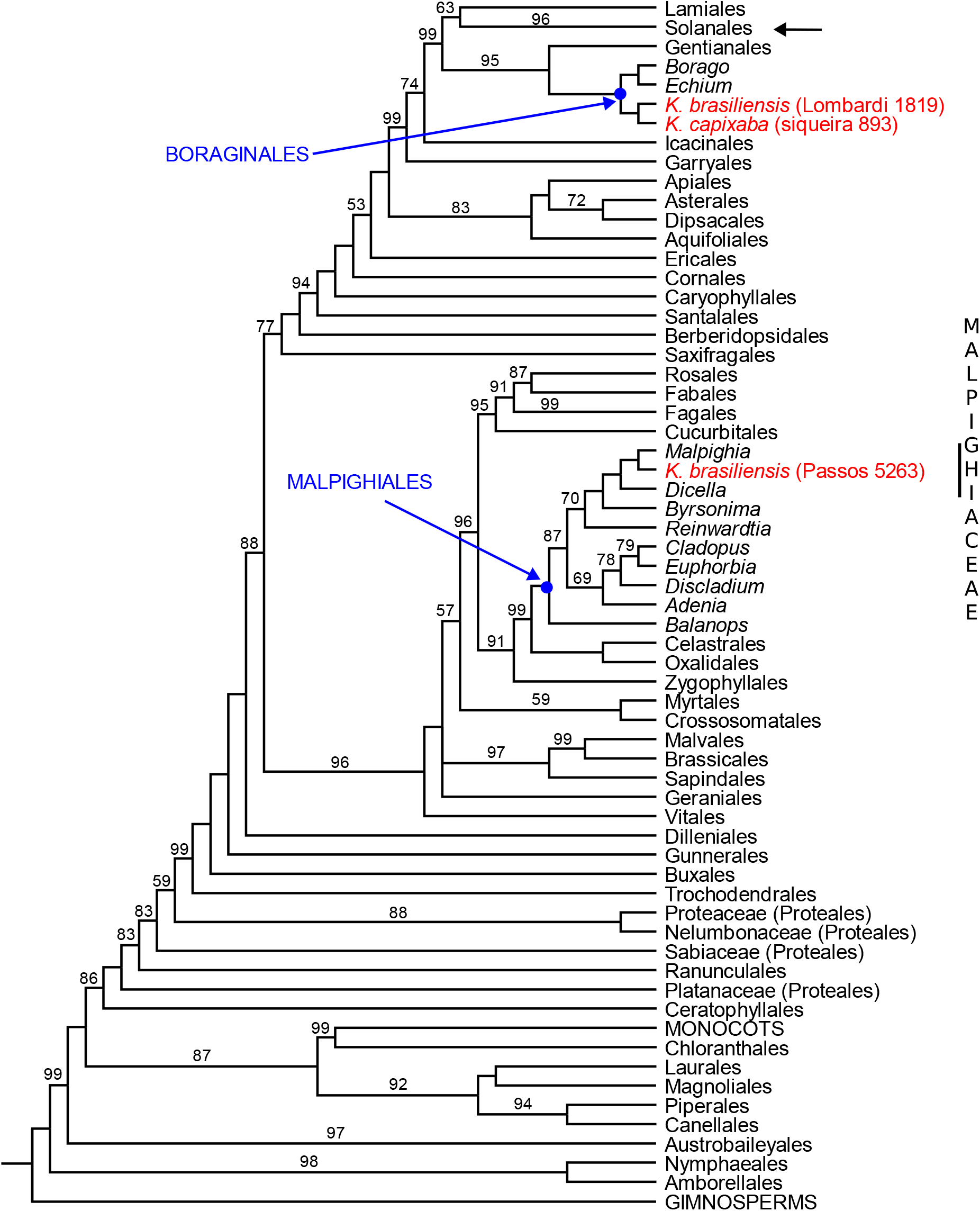
Summary Maximum Likelihood Angiosperm phylogeny showing the position of the three specimens newly sequenced in this study (in red). Phylogeny inferred using the chloroplast *matK* region; numbers on the branches represent IQ-Tree Ultrafast Bootstrap support values; branches without numbers received 100% support. Passos 5263 (*Keraunea brasiliensis*) is nested in Malpighiales; Lombardi 1819 (*K. brasiliensis*) and Siqueira 893 (*K. capixaba*) are nested in Boraginales. The blue arrows indicate the position of Malpighiales and Boraginales; the black arrow indicates the position of Solanales, where the family Convolvulaceae belongs. See complete phylogeny in Supplementary File 2.

Both BLAST and phylogenetic analysis indicate the two *Keraunea brasiliensis* specimens and the one *K. capixaba* specimen sequenced are not closely related to each other, and that *K. brasiliensis* is nested within Malpighiaceae Juss. (order Malpighiales) and *K. capixaba* is nested within Ehretiaceae Mart. (order Boraginales. In other words, the genus *Keraunea* is polyphyletic with its two constituent species belonging to Malpighiaceae and Ehretiaceae rather than Convolvulaceae as originally classified. The sequences obtained were of high quality and showed no evidence of contamination, as would be expected since the labs where the DNA was processed had not sequenced any material of these families. We subsequently inferred densely-sampled phylogenies of the two families where these specimens were nested, Malpighiaceae and Ehretiaceae, and the results for each specimen are described below.

### *L. Passos 5263* specimen, the isotype of *Keraunea brasiliensis*

All three regions (*matK, rbcL* and *nrITS*) amplified for *Passos 5263*, the isotype of *Keraunea brasiliensis*, indicate it is nested within the family Malpighiaceae. To further explore its position within the family, we inferred three species-level Malpighiaceae phylogenies — *matK* (TVM+F+G4), *matK+rbcL* (TVM+F+I+G4), and *nrITS* (TIM2+F+I+G)— with GenBank data and including representatives of all genera accepted in the most recent review of the family [18] (Supplementary File 1). We used several *Bergia* L. (Elatinaceae Dumort), *Elatine* L. (Elatinaceae), *Bhesa* (Centroplacaceae Doweld & Reveal) and *Euphorbia maculata* L. (Euphorbiaceae Juss.) specimens as outgroups following [19–21], with *Euphorbia maculata* used to root the phylogenies. The *matK* phylogeny places *Passos 5263* in a clade with *Mascagnia affinis* W.R.Anderson & C.Davis, *M. cordifolia* (A.Juss.) Griseb., and *M. dissimilis* C.V.Morton & Moldenke, all three species also present in Brazil^1^ (Figure 2a) and corresponding to the core/true *Mascagnia* rather than the many recent segregates of this former polyphyletic genus [22–26]. This clade of three species is also retrieved, with higher support, in [18], a phylogeny based on *matK* and *rbcL* plus two additional regions not sequenced in our study, chloroplast *ndhF* and nuclear *PHYC*.

**Figure 2.**
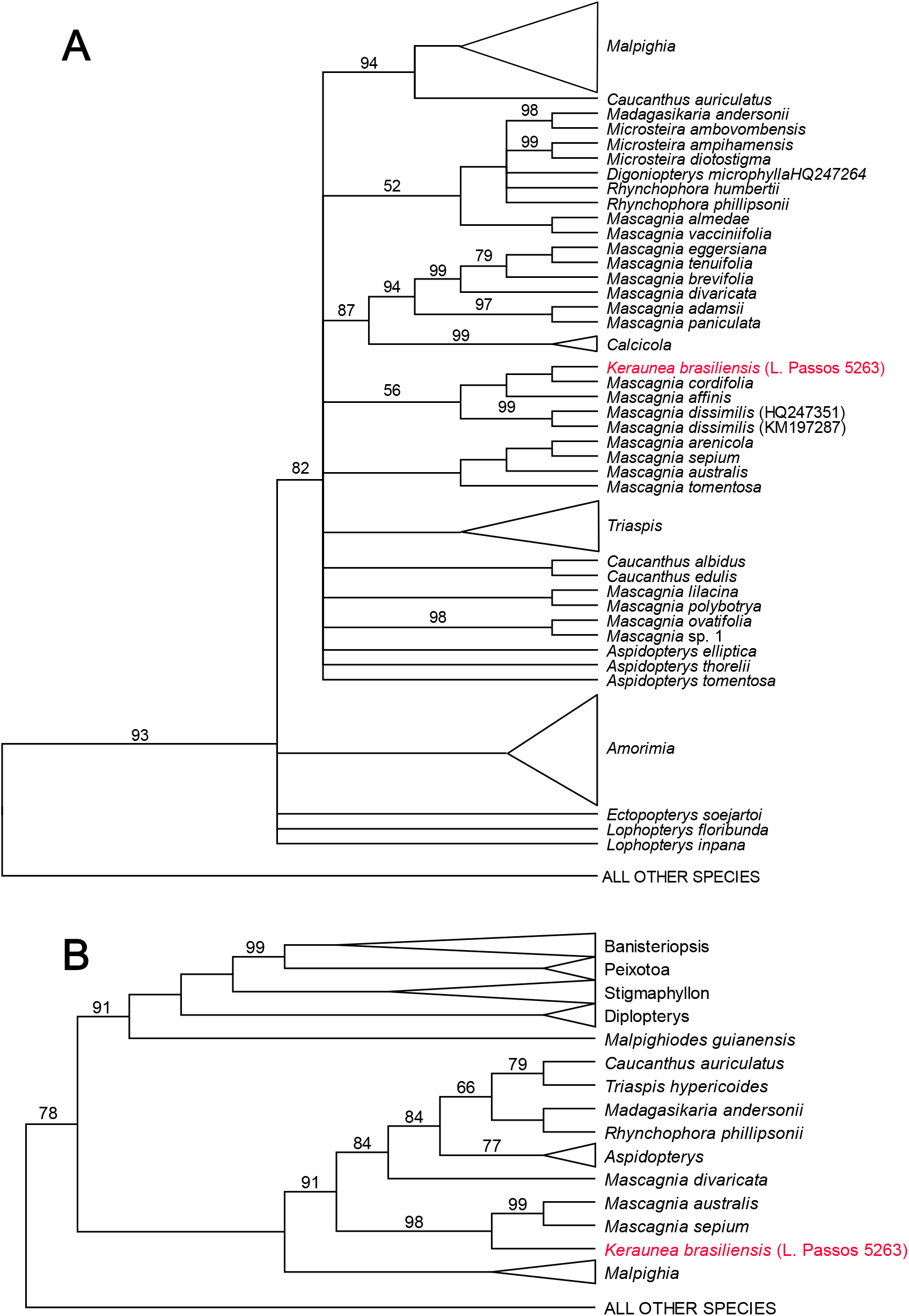
Summary Maximum Likelihood Malpighiaceae phylogenies showing the position of the *Keraunea brasiliensis* Passos 5263 specimen (in red). Phylogenies inferred using (A) the chloroplast *matK* region and (B) the *nrITS* region; numbers on the branches represent IQ-Tree Ultrafast Bootstrap support values; branches without numbers received 100% support. Passos 5263 (Keraunea brasiliensis) is nested in Mascagnia in both phylogenies. See complete phylogenies in Supplementary File 3.

The *nrITS* phylogeny (Figure 2b) places *Passos 5263* in a clade with three *Mascagnia* species widely distributed in South America: *M. australis* C.E.Anderson, *M. divaricata* (Kunth) Nied. and *M. sepium* (A.Juss.) Griseb. Again, *M. australis* and *M. divaricata* are within the same *Mascagnia* clade in [18], whereas *M. sepium* was not included in that study but is also taxonomically recognised as a member of core/true *Mascagnia*.

In summary, our molecular results strongly suggest *Passos 5263* is a Malpighiaceae, most likely a *Mascagnia*, and would therefore justify transferring the species *Keraunea brasiliensis —*and therefore the genus *Keraunea pro parte*—, to this family. However, these molecular results are particularly surprising because *K. brasiliensis* does not exhibit the canonical ‘Malpighiaceae morphology’. Members of Malpighiaceae usually present simple, opposite leaves with T-shaped, unicellular trichomes and often inter-petiolar stipules, oil glands present on the sepals and/or extra-floral glands on the petiole or leaf blade, and five free, usually clawed petals [27,28]. In contrast, leaves in *Passos 5263* seem to be alternate, and oil glands cannot be readily observed on this specimen’s sepals. Moreover, this specimen seems to have a sympetalous perianth, the branching of the inflorescence is alternate, and the fruit is very peculiar if Malpighiaceae. In addition, although the molecular results reported here are robust, considering the strong morphological differences between *K. brasiliensis* and the members of Malpighiaceae we hesitate to formally propose this taxonomic change. We think further study is advisable before *Keraunea brasiliensis* can be re-classified. An additional trait to explore is the three carpellate ovary with a single locule per ovule as is found in Malpighiaceae. Finally, we have not seen the holotype (deposited at Sao Paulo herbarium (SP) and not yet digitised) and it may be different in whole or in part from the isotype at Kew. In conclusion, it seems clear that *Keraunea brasiliensis* is not a Convolvulaceae and future studies should be able to determine its generic placement within Malpighiaceae.

### *J. A. Lombardi 1819* and *G. Siqueira 893* specimens

We also sequenced a *Keraunea brasiliensis* paratype —*Lombardi 1819* (K)— and a *K. capixaba* paratype —*Siqueira 893* (K)—. We sequenced *matK* and *nrITS* for *Lombardi 1819* and *matK* for *Siqueira 893*. We first checked our sequences against the GenBank database using BLAST, as well as incorporated the *matK* sequences to the Angiosperm phylogenies aforementioned (Figure 1). Interestingly, these two paratype specimens are nested in Boraginales and appear to be closely related to *Ehretia* P.Browne, *Halgania* Gaudich. and other genera in the family Ehretiaceae, recently segregated from Boraginaceae Juss. [29].This was confirmed with a *matK* (TVM+F+G4) phylogeny of all accepted genera in Ehretiaceae [cf. 30,31] and the other families in the order (Figure 3a). In this *matK* phylogeny, both *Keraunea* specimens (*Lombardi 1819* and *Siqueira 893*) are most closely related to each other, and sister to a monophyletic genus *Ehretia*, with high support. The close relationship between *Lombardi 1819* and *Siqueira 893* is not surprising as these two specimens are remarkably similar morphologically.

**Figure 3.**
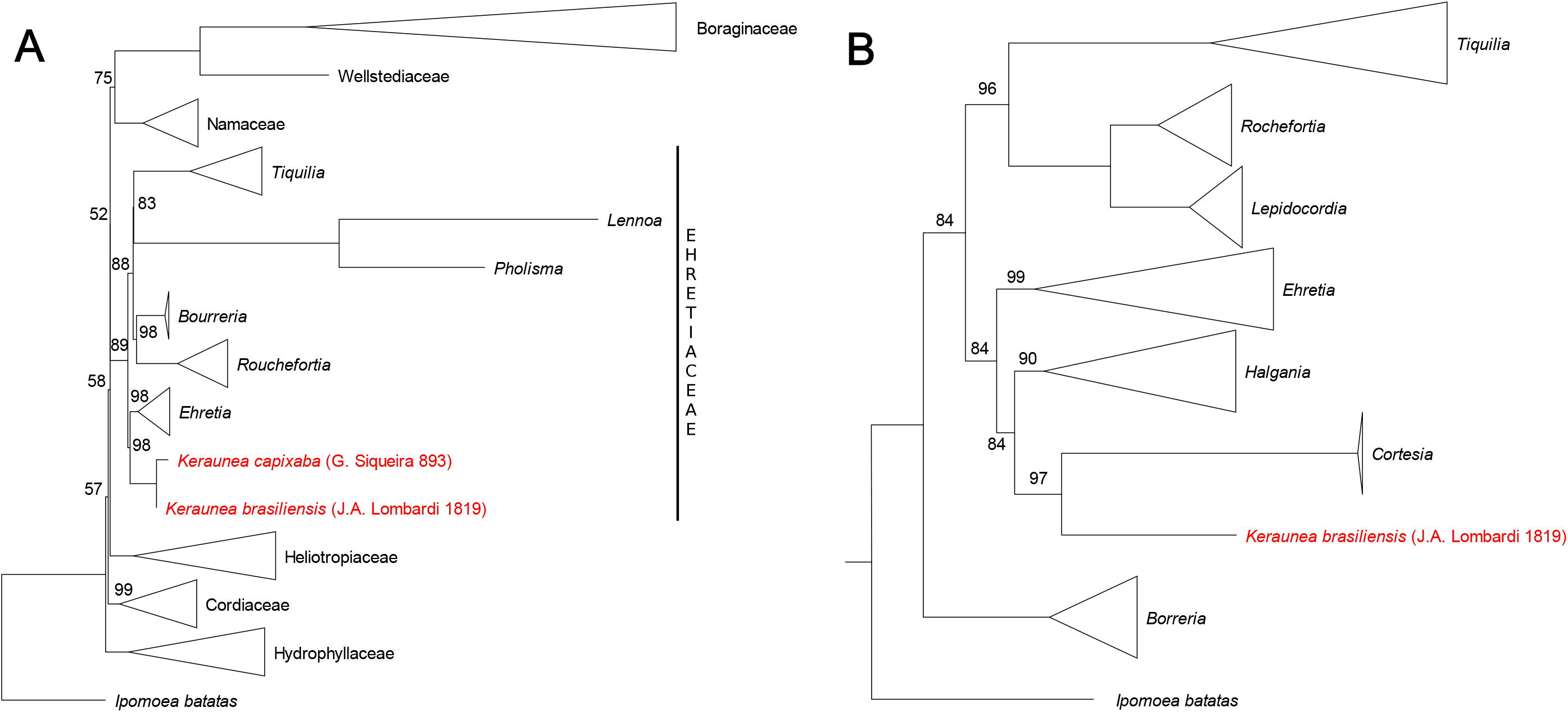
Summary Maximum Likelihood phylogenies showing the position of *Keraunea brasiliensis* specimen *J.A. Lombardi 1819* and *K. capixaba* specimen *G. Siqueira 893* in Ehretiaceae. (A) Boraginales phylogeny inferred using the chloroplast *matK* region. (B) Ehretiaceae phylogeny inferred using the *nrITS* regions. Triangles represent monophyletic family and/or genera with multiple specimens; numbers on the branches represent IQ-Tree Ultrafast Bootstrap support values; branches without numbers received 100% support. In (A), the two specimens newly sequenced in this study, *Lombardi 1819* and *Siqueira 893*, are most closely related to each other and sister to a monophyletic genus *Ehretia*, with high support. In (B), *Lombardi 1819* is most closely related to a monophyletic genus *Cortesia*, which is not included in (A). See complete phylogenies in Supplementary file 4.

A densely sampled *nrITS* phylogeny of Ehretiaceae (GTR+F+I+G4) places *Lombardi 1819* sister to monophyletic *Cortesia cuneifolia* Cav. (= *Ehretia cortesia* Gottschling), a genus recently segregated by Gottschling *et al*. [31]. This clade is sister to a monophyletic *Halgania*, also with moderately high support (Figure 3b). It is important to note that *Cortesia* Cav. and *Halgania* have not been sequenced for *matK* and therefore they are not included in the *matK* phylogeny.

The results of both *nrITS* and *matK* phylogenies presented here are congruent and show the two *Keraunea* collections *Lombardi 1819* and *Siqueira 893* are most closely related to *Cortesia, Halgania* and *Ehretia*. Again, although our molecular results are robust, we think a comprehensive, complete study of *Keraunea* is needed before *Keraunea capixaba* and at least one of the *K. brasiliensis* paratypes can be re-classified. It seems clear that *K. capixaba* is also not a Convolvulaceae, and future studies should be able to confirm its placement in the right family, most likely Ehretiaceae.

## CONCLUSIONS

Here, we have shown that *Keraunea brasiliensis* and *K. capixaba* comprise a mixture of specimens from different Angiosperm families not Convolvulaceae. We have refrained from proposing formal taxonomic changes as we think a comprehensive study of all material assigned to *Keraunea* is necessary, including the broader context of the respective families where the two species belong. It is unlikely that the mere addition of more molecular data will clarify the position of these two species, at least for their family designations, unless such a study is accompanied by a comprehensive taxonomic study of herbarium and living material. The results of our study highlight several important aspects of contemporary plant systematics. First, the inadequate role of gross morphology in placing some material in the correct family. Second, the ability of molecular sequence data to quickly place difficult specimens in the correct family. Third, the large fraction of taxa that still remain to be classified and fully understood.

## Footnotes

1 Adding the chloroplast *rbcL* region slightly improves phylogenetic resolution, but the only two *Mascagnia* species (*M. adamsii* and *M. sepium*) that have data available for both *matK* and *rbcL* seem to be only distantly related to *Passos 5263*. See Supplementary File 3.

